# CPT2 mediated fatty acid oxidation is dispensable for humoral immunity

**DOI:** 10.1101/2024.05.15.594133

**Authors:** Meilu Li, Xian Zhou, Xingxing Zhu, Yanfeng Li, Taro Hitosugi, Yuzhen Li, Hu Zeng

**Affiliations:** Division of Rheumatology, Department of Medicine, Mayo Clinic Rochester, MN 55905, USA; Department of Dermatology, The Second Affiliated Hospital of Harbin Medical University, Harbin, 150001, P. R. China; Department of Oncology, Mayo Clinic, Rochester, MN 55905, USA; Department of Immunology, Mayo Clinic Rochester, MN 55905, USA

## Abstract

B cell activation is accompanied by dynamic metabolic reprogramming, supported by a multitude of nutrients that include glucose, amino acids and fatty acids. While several studies have indicated that fatty acid mitochondrial oxidation is critical for immune cell functions, contradictory findings have been reported. Carnitine palmitoyltransferase II (CPT2) is a critical enzyme for long-chain fatty acid oxidation in mitochondria. Here, we test the requirement of CPT2 for humoral immunity using a mouse model with a lymphocyte specific deletion of CPT2. Stable ^13^C isotope tracing reveals highly reduced fatty acid-derived citrate production in CPT2 deficient B cells. Yet, CPT2 deficiency has no significant impact on B cell development, B cell activation, germinal center formation, and antibody production upon either thymus-dependent or –independent antigen challenges. Together, our findings indicate that CPT2 mediated fatty acid oxidation is dispensable for humoral immunity, highlighting the metabolic flexibility of lymphocytes.

## INTRODUCTION

In response to challenges from thymus-independent (TI) or thymus-dependent (TD) antigens, B cells can be activated and differentiate into plasmablasts or form germinal centers (GCs), which generate memory B cells and long-lived plasma cells (1–3). Plasmablasts and plasma cells produce antibodies to fight against pathogen infection. B cells are also a major source of inflammatory cytokines. Thus, B cells are vital for host defense and the success of vaccines.

B cell differentiation is accompanied by dynamic metabolic reprogramming (4). For example, glucose metabolism is critical for B cell development, activation, GC formation, and transformation (5–8). Furthermore, mitochondrial oxidative phosphorylation is critical for B cell survival and differentiation (9–11). Beyond glucose, fatty acids are a major energy source (12). Long chain fatty acid (LCFA) beta-oxidation requires transportation of LCFA across mitochondrial membranes, which is carried out by carnitine palmitoyltransferase I (CPT1) and II (CPT2). CPT1 has two isoforms, CPT1A and CPT1B. Deletion of CPT1A or CPT2 prevents LCFA beta-oxidation (13, 14). Although LCFAs could in theory contribute to mitochondrial oxidative phosphorylation, the importance of CPT-mediated LCFA oxidation for immune cells remains contentious. Previous studies showed that the differentiation of T cells and macrophages is not affected by ablation of CPT1A or CPT2 (15, 16). In contrast, NK cell proliferation, anti- viral and anti-tumor effector functions are compromised without CPT1A, indicating that the dependency of CPT-mediated LCFA oxidation could be cell type specific. Furthermore, GC B cell formation is partially suppressed when CPT2 expression is reduced through shRNA-mediated gene knockdown in B cells, although whether *Cpt2* knock-down impairs antibody production remains unknown (17). To further address the role of fatty acid oxidation in B cell function, we assessed humoral immunity in a mouse line with lymphocyte specific deletion of *Cpt2* and bone marrow chimeric mice with B cell specific ablation of *Cpt2*. Our data demonstrate that CPT2 is required for optimal LCFA beta-oxidation in B cells, but it is dispensable for B cell activation *in vitro* and humoral immunity *in vivo*. These results indicate that LCFAs may influence B cell functions independent of fatty acid oxidation in mitochondria.

## MATERIALS AND METHODS

### Mice

*Cd2^iCre^Cpt2*^fl/fl^ mice were generated by crossing *Cpt2*^fl/fl^ mice with *Cd2-iCre* transgenic mice (Jackson Laboratory). C57BL/6, B6.CD45.1 (B6.SJL-*Ptprc^a^ Pepc^b^*/BoyJ), and *Rag1*^−/−^ mice were purchased from the Jackson Laboratory. Chimeric mice model was generated by transferring 2.5 × 10^6^ bone marrow (BM) cells isolated from *Cd2^iCre^Cpt2*^fl/fl^ or *Cpt2*^fl/fl^ (wild type, WT) depleted T cells, mixed with 2.5 × 10^6^ BM cells from B6.CD45.1 mice into *Rag1*^−/−^ mice. After 4 weeks, mice were immunized intraperitoneally with NP-OVA/Alum. Mice were maintained under pathogen-free conditions and were treated in accordance with the guidelines of the Department of Comparative Medicine of Mayo Clinic with approval from the Institutional Animal Care and Use Committee (IACUC).

Preparation for NP-OVA/Alum immunization by mixing 100 µg NP-OVA (BioSearch Technologies) and 10% KAL(SO_4_)_2_ (Sigma) dissolved in PBS at a ratio of 1:1, together with 10 µg LPS (Sigma-Aldrich) at pH 7. For TNP-LPS immunization, 50 µg of TNP-LPS (Biosearch Technologies) was dissolved in 200 µl PBS and administered to each mouse via intraperitoneal injection.

### Influenza Virus Infection

For influenza virus infection, influenza A/PR8 strain (60 pfu/mouse) were diluted in FBS-free DMEM media on ice and inoculated in anesthetized mice through intranasal route. Sera were collected before and two weeks after infection for fatty acid component measurement. The mediastinal lymph nodes and spleens were harvested and analyzed for germinal center B cell and plasmablast formation and follicular helper T differentiation.

### Immune cell purification and culture

Mouse B cells were isolated from spleen single cell suspension using Mouse B Cell Isolation Kit (StemCell Technologies). B cells were cultured in RPMI 1640 (Corning) containing 10% FBS (Gibco), 10 mM HEPES (Gibco), Penicillin Streptomycin Glutamine mixed solution (Gibco) and 50 µM 2-Mercaptoethanol (Sigma-Aldrich), stimulated with 10 µg/mL LPS (Sigma-Aldrich), 10 ng/mL recombinant IL-4 (Tonbo Bioscience) and 20 ng/mL recombinant BAFF (Biolegend). B cells were activated with 2.5 µM TLR ligand CpG (InvivoGen), 10 ng/mL recombinant IL-4 and 10 ng/mL IL-5 (Tonbo Bioscience). B cells were treated with or without SSI-4 (gift from John A. Copland, III, Mayo Clinic, Florida, USA) or Oleic Acid-Albumin from bovine serum (Sigma-Aldrich) or Thioridazine (Cayman chemical). B cell proliferation was measured by dilution of CellTrace Violet proliferation dye (Life Technologies).

### Real-time PCR

For mRNA analysis, total mRNA was extracted from mouse B cells by RNeasy Micro kit (QIAGEN) and converted to cDNA by using PrimeScript RT Master Mix (Takara), following the manufacturer’s protocols. This was done in preparation for subsequent real-time PCR analysis by using a Takara Realtime PCR system. The following primers were used, *Cpt1a* primers, forward, 5’-CTCCGCCTGAGCCATGAAG-3’, reverse, 5’-CACCAGTGATGATGCCATTCT-3’; *Cpt1b* primers, forward, 5’- GCACACCAGGCAGTAGCTTT-3’, reverse, 5’- CAGGAGTTGATTCCAGACAGGTA-3’; *Cpt2* primers, forward, 5’- GGATTTTGAGAACGGCATTGG-3’, reverse, 5’-TTAAAACGACAGAGTCTCGAGC-3’. β-actin expression was used as control.

### Metabolic Assays

The bioenergetic activities were measured using a Seahorse XFe96 Extracellular Flux Analyser following established protocols (Agilent). Briefly, about 300,000 B cells were seeded per well on Cell-Tak (Corning) coated XFe96 plate with fresh XF media (Seahorse XF RPMI medium containing 10 mM glucose, 2 mM L-glutamine, and 1 mM sodium pyruvate, PH 7.4; all reagents from Agilent). Oxygen consumption rates (OCR) were measured in the presence of Oligomycin (1.5 µM; Sigma-Aldrich), FCCP (1.5 µM; Sigma-Aldrich), and Rotenone (1 µM; Sigma-Aldrich)/ Antimycin A (1 µM; Sigma-Aldrich) in Mito Stress assay.

### Flow Cytometry

Single-cell suspension from spleens, peripheral lymph nodes, thymus, bone marrow or Payer’s patches or mediastinal lymph nodes were prepared in PBS containing 2% (w/v) FBS on ice. For analysis of surface markers, cells were stained in PBS containing 1% (w/v) BSA on ice for 30 min, with BV605-labelled B220 (clone: RA3-6B2; BioLegend), APC-labelled anti-CD43 (clone: 53-6.7; BD), Percp-cy5.5 -labelled anti-CD19 (clone: 1D3; Tonbo), PE-Cy7-labelled anti-CD21 (clone: IM7; BioLegend), FITC-labelled anti-CD23 (clone: B3B4; BioLegend), Alexa Fluor647- labelled anti-CD95/Fas (clone: Jo2; BD Biosciences), Pacific Blue-labelled anti-GL7 (clone: GL7; BioLegend), APC-Cy7-labelled TCRβ (clone: H57-597; BioLegend), BV605-labelled anti- CD4 (clone: RM4-5; BioLegend), PE-labelled anti-CD8 (clone: 53-6.7; BioLegend), FITC-labelled anti-IgG1 (clone: RMG1-1; BioLegend), BV711-labelled anti-CD138 (clone: PC61; BioLegend), APC/Cy7-labelled anti-IgD (clone: 11-26c.2a; BioLegend), APC-labelled anti-ICOS (clone: 7E.17G9; BD Biosciences), PE-Cy7-labelled anti PD-1 (clone: RMPI-30; BioLegend), BV605-labelled anti-CD25 (clone: 281-2; BioLegend), PE-Cy7-labelled anti-CD24 (clone: M1/69; BioLegend), BV711 -labelled anti-CD44 (clone: IM7; BioLegend). Cell viability was stained using the Ghost Dye Violet 510 (Cytek). CXCR5 was stained with biotinylated anti- CXCR5 (clone: 2G8, BD Biosciences) followed by staining with streptavidin-conjugated PE (BD Biosciences). Transcriptional factors FoxP3 (clone: FJK-16 s; eBioscience) was stained using the Transcription Factor Buffer Set (True-Nuclear, BioLegend). Flow cytometry was performed on the Attune NxT (ThermoFisher) cytometer, and analysis was performed using FlowJo software (BD Biosciences).

### Mouse Immunoglobulin Isotyping Panel Detection

To detect the concentrations of mouse immunoglobulin isotypes IgG1, IgG2a, IgG2b, IgG3 and total IgM and IgA in sera, we used LEGENDplex mouse immunoglobulin isotyping panel, according to manufacturer’s instructions (Biolegend).

### ELISA

To measure the levels of NP-specific antibodies in sera, 96-well plates (2596; Costar) were coated with 1 µg/mL NP23-BSA or NP2-BSA (Biosearch Technologies) in PBS and incubated at 4°C overnight. The plates were washed four times using 0.05% Tween 20 in PBS, then blocked with 5% blocking protein (Bio-Rad) for 1.5h at 37°C and washed four times again. Serum samples, serially diluted with 1% BSA, were incubated in the plates for 1.5h at 37°C and then washed eight times. Horseradish peroxidase (HRP)-conjugated detection antibodies for total IgG, IgG1, IgG2c, and IgM (Bethyl Laboratories) were added and incubated at room temperature for 1h, followed by eight washes. The reaction was developed with tetramethylbenzidine (TMB) substrate and stopped with 2N H_2_SO_4_. The absorbance was read at a wavelength of 450 nm. Similar methods were used to detect antibodies specific to the influenza A/PR8 strain (a gift from Dr. Jie Sun, University of Virginia, USA) and TNP-LPS (Biosearch Technologies) in sera.

### Mass Spec Based-FAO Activity Assay

FAO activity was measured by monitoring the conversion rate of [U-^13^C]-palmitate to [M+2, 4, 6]-labeled ^13^C citrate with GC/MS. In brief, after 48 h activation with LPS/IL-4/Baff, the cells were incubated with 100 mM [U-^13^C]-palmitate-BSA conjugate for overnight. After the incubation, the cells were quickly rinsed in ice-cold PBS and lysed with 80% methanol. The crude lysates were centrifuged to remove debris. The resulting supernatants were dried with nitrogen gas, dissolved in 75 μL DMF, then derivatized with 75 μL N-Methyl-N-(tert-Butyldimethylsilyl) trifluoroacetamide (MTBSTFA)+ 1% tert-Butyldimetheylchlorosilane (TBDMCS) (Regis). Samples were incubated at room temperature for 30 minutes and were analyzed using an Agilent 7890B GC coupled to a 5977A mass detector. 3 μL of derivatized sample were injected into an Agilent HP-5ms Ultra Inert column, and the GC oven temperature increased at 15°C/min up to 215°C, followed by 5°C/min up to 260°C, and finally at 25°C/min up to 325°C. The MS was operated in split-less mode with electron impact mode at 70 eV. Mass range of 50-700 was analyzed, recorded at 1,562 mass units/second. Data was analyzed using Agilent MassHunter Workstation Analysis and Agilent MSD ChemStation Data Analysis software. IsoPat2 software was used to adjust for natural abundance as previously performed (18). The extent of isotopic ^13^C labeling in citrate was further divided by percent isotopic enrichment of intracellular [U-^13^C]-palmitate to determine the conversion rate of [U-^13^C]- palmitate to [M+2, 4, 6]-labeled citrate in cells. Palmitate and citrate were detected by GC-MS as TBDMS derivatives at the following m/z values: palmitate (m/z 313), [U-^13^C]-palmitate (m/z 329), and citrate (m/z 459-465).

### Statistical Analysis

Statistical analysis was performed using GraphPad Prism (version 9). Comparisons between two groups were examined using a two-tailed Student’s t-test. One-way ANOVA or two-way ANOVA was performed for comparisons involving more than three groups. A p-value < 0.05 was considered statistically significant.

## RESULTS

### The absence of CPT2 does not substantially impact the development or homeostasis of B and T cells

To study the function of CPT2 in adaptive immunity, we crossed a floxed *Cpt2* allele with *Cd2- iCre* transgenic mice to generate *Cd2^iCre^Cpt2*^fl/fl^ mice, in which *Cpt2* is efficiently deleted in lymphocyte lineages (19, 20). Quantitative Real-time PCR confirmed that *Cpt2* was efficiently ablated in B cells from *Cd2^iCre^Cpt2*^fl/fl^ mice, while expression of *Cpt1a* and *Cpt1b* remained normal (Figure 1A). Next, we examined the development of B and T cells. *Cd2^iCre^Cpt2*^fl/fl^ mice harbored normal percentages and number of B220^int^CD43^+^ B cell precursors, B220^int^CD43^low^ pre-B/immature B cells, B220^hi^CD43^low^ mature B cells, and CD138^+^ plasma cells in the bone marrow (BM) compared to WT mice (Figure 1B, 1C). In the spleen, WT and *Cd2^iCre^Cpt2*^fl/fl^ mice had equivalent numbers of CD21^int^CD23^+^ follicular (Fo) and CD21^hi^CD23^low^ marginal zone (MZ) B cells (Figure 1D). The spontaneous formation of GL7^+^Fas^+^ germinal center (GC) B cells in Peyer’s patches was not affected by CPT2 deficiency (Figure 1E). To examine T cell development and homeostasis, we quantified T cells subsets in thymus and spleens. Thymus from *Cd2^iCre^Cpt2*^fl/fl^ mice had normal percentages of CD4^−^CD8^−^ double negative (DN), CD4^+^CD8^+^ double positive (DP), CD4^+^ and CD8^+^ single positive (SP) cells. Although the absolute number of DP thymocytes was modestly and significantly reduced in KO mice, the number of CD4^+^ and CD8^+^ SP thymocytes were normal, suggesting that CPT2 deficiency reduces DP thymocyte formation but does not affect the final maturation of SP thymocytes (Figure 1F). Consistent with this observation, *Cd2^iCre^Cpt2*^fl/fl^ mice had normal percentages and numbers of CD4^+^, CD8^+^ T cells, and CD4^+^Foxp3^+^ regulatory T cells (Tregs) in the spleens (Figure 1G). Therefore, these data indicate that CPT2 is largely dispensable for B and T cell development and homeostasis.

**FIGURE 1.**
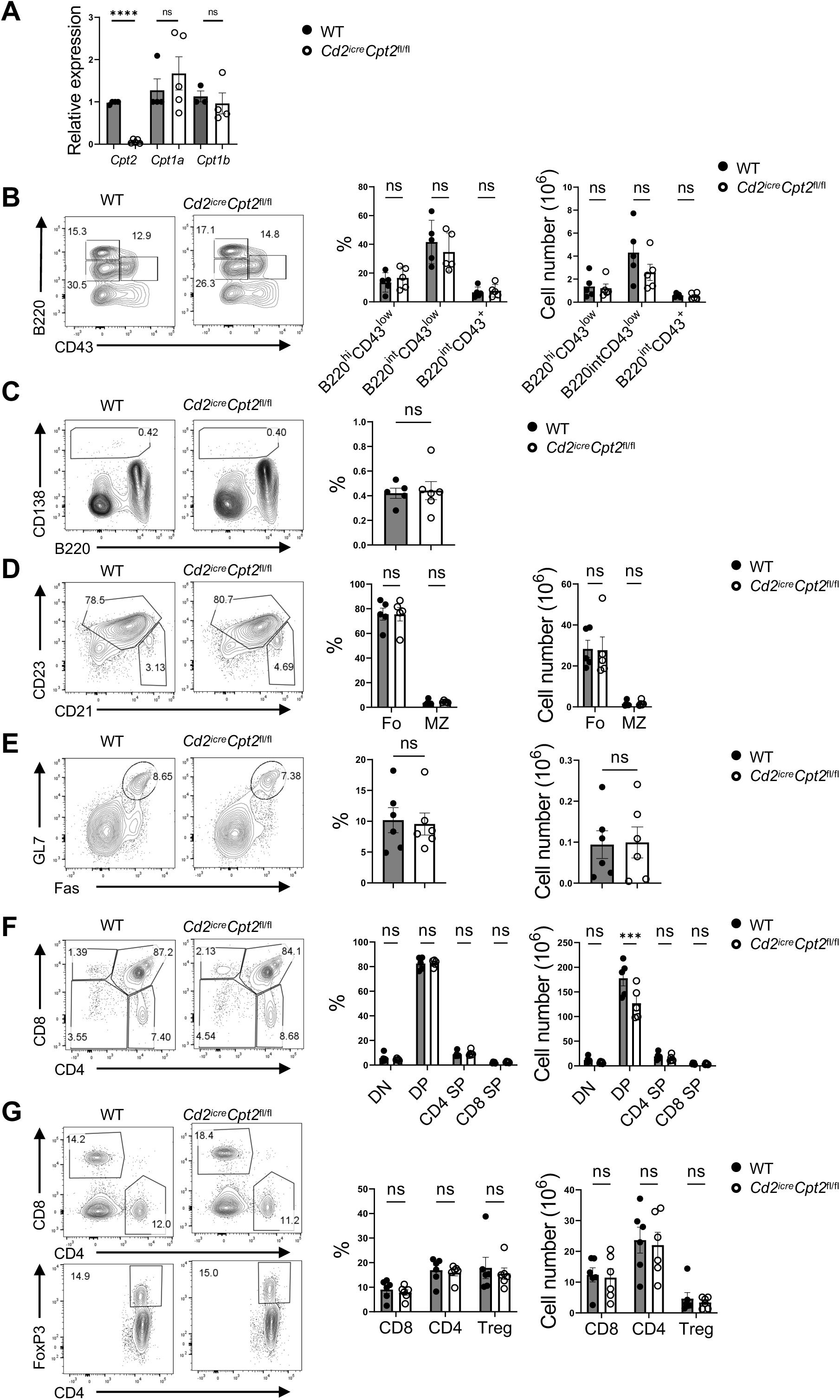
*Cd2^iCre^Cpt2*^fl/fl^ mice have largely normal B and T cell development and homeostasis. (A) Real-time PCR analysis of *Cpt2*, *Cpt1a* and *Cpt1b* from B cells isolated from spleens. The expression level of *Cpt2* in WT mice was used as an internal control for RNAs. (B) Left, the expression of B220 and CD43 among BM cells analyzed by flow cytometry. Right, the frequency (left) and absolute number (right) of B220^hi^CD43^low^, B220^int^CD43^low^ and B220^int^CD43^+^ cell populations. (C) Left, the expression of B220 and CD138 among BM cells analyzed by flow cytometry. Right, the frequency of CD138^+^ populations. (D) Left, flow cytometry analysis of Fo (CD21^low^CD23^+^), MZ (CD21^hi^CD23^−^) cells among B220^+^CD19^+^ population from spleens. Right, the frequency (left) and absolute number (right) of Fo and MZ cell populations. (E) Left, flow cytometry analysis of GC (GL7^+^Fas^+^) among TCRβ^-^CD19^+^ population from Peyer’s patches. Right, the frequency (left) and absolute number (right) of GC populations. (F) Left, the expression of CD4 and CD8 among thymus lymphocytes analyzed by flow cytometry. Right, the frequency (left) and absolute number (right) of DN (CD4^-^CD8^-^), DP (CD4^+^CD8^-^), CD4 SP (CD4^+^CD8^-^) and CD8 SP (CD4^-^CD8^+^) cell populations. (G) Left, flow cytometry analysis of CD4, CD8 and Treg cells (B220^-^CD4^+^FoxP3^+^) from spleens. Right, the frequency (left) and absolute number (right) of CD4, CD8 and Treg cells. (A-G) Cells were from WT and *Cd2^iCre^ Cpt2*^fl/fl^ mice. *p* values were calculated using two-way ANOVA (A, B, D-G) and Student’s t-test (C). ns, not significant; \*\*\**p* < 0.001. Data were collected from at least 3 (A-G) independent experiments. Error bars represent SEM.

### CPT2 deficiency reduces fatty acid oxidation and oxidative phosphorylation in B cells

Because CPT2 is obligatory for LCFAs to be transported into the mitochondrial matrix for beta- oxidation, we predicted that *Cpt2* deletion would reduce palmitate oxidation in B cells. To this end, we assessed FAO activity in B cells from spleens of WT and *Cd2^iCre^Cpt2*^fl/fl^ mice by monitoring the conversion rate of [U-^13^C]-palmitate to [M+2, 4, 6]-labeled ^13^C citrate using gas chromatography coupled with mass spectrometry. CPT2 deficiency significantly reduced the [U- ^13^C]-palmitate oxidation rate (Figure 2A), demonstrating that CPT2 is required for efficient LCFA beta-oxidation in B cells. In agreement with this phenotype, CPT2 deficient B cells exhibited modest, but significant, reduction of basal respiration, ATP linked respiration, maximal respiration and spared respiratory capacity measured by Seahorse bioanalyzer compared to WT B cells (Figure 2B). Our previous study showed that monounsaturated fatty acids, such as oleic acid (OA), can promote B cell mitochondrial metabolism (21). To our surprise, OA promoted basal respiration and ATP linked respiration in both WT and CPT2 deficient B cells (Figure 2B). These data indicate that CPT2 is necessary for optimal mitochondrial oxidative phosphorylation, but the respiration boosting effect of exogenous OA is independent of CPT2. In another word, our results indicate that LCFA could improve B cell mitochondrial metabolism without undergoing CPT2-mediated LCFA beta-oxidation.

**FIGURE 2.**
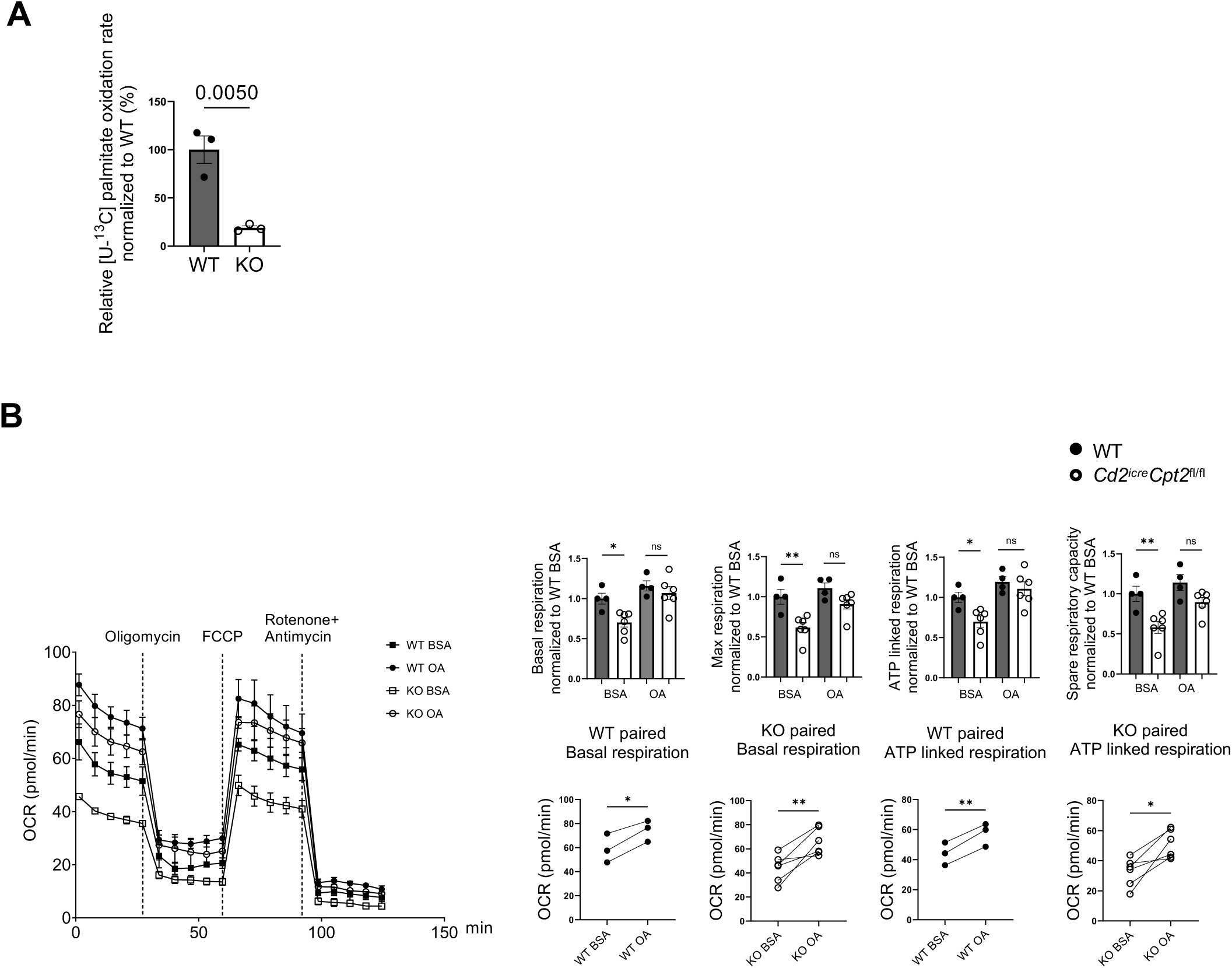
*Cpt2* deficiency leads to a defect in FAO in B cells. (A) FAO activity was measured by [U-^13^C] palmitate oxidation rate in B cells from WT and *Cd2^iCre^Cpt2*^fl/fl^ mice. (B) Oxygen consumption rate (OCR) was measured using a Seahorse XFe96 analyzer using CpG/IL4/IL5 activated B cells treated with BSA (50 µM) as control, or OA (100 µM) for 48 h with MitoStress Test. Right: summary of basal respiration, ATP-linked respiration, max respiration, and spare respiratory capacity. Data are representative of at least three independent experiments. *p* values were calculated using Student’s t-test (A-B). ns, not significant, \**p* < 0.05, \*\**p* < 0.01. Error bars represent SEM.

### CPT2 is dispensable for B cell activation and proliferation *in vitro*

Next, we sought to determine whether *Cpt2* ablation affected B cell activation *in vitro*. We labeled WT and KO B cells with CellTrace violet (CTV) and stimulated the cells with lipopolysaccharide (LPS)/IL-4/Baff for 3 days. Flow cytometry analysis showed that WT and CPT2 deficient B cells had equivalent CTV dilution, indicating similar cell division (Figure 3A). The inhibitor of stearoyl-CoA desaturase (SCD), SSI-4, strongly suppressed B cell proliferation, which was rectified by exogenous OA in both WT and CPT2 KO B cells (Figure 3A). Similar observation was observed in terms of IgG1 class switch (Figure 3B), suggesting that SCD inhibition might not impact CPT2-mediated FAO. Exogenous OA could enhance IgG1 class switch in WT and CPT2 deficient B cells stimulated with LPS/IL-4/Baff (Figure 3C). Such enhancement was not limited to TLR4 engagement. OA could promote IgG1 expression upon stimulation with toll-like receptor 9 (TLR9) ligand CpG/IL-4/IL-5 in the presence or absence of CPT2, with CPT2 deficient B cells exhibiting even higher IgG1 expression than WT B cells (Figure 3D). FAO could take place in the mitochondria as well as peroxisome (22). Therefore, we hypothesized that peroxisomal FAO may compensate for the loss of CPT2. We treated B cells with thioridazine, a peroxisome inhibitor, and observed that both WT and CPT2 null B cells exhibited reduced proliferation and class switching upon thioridazine treatment (Figure 3E).

**FIGURE 3.**
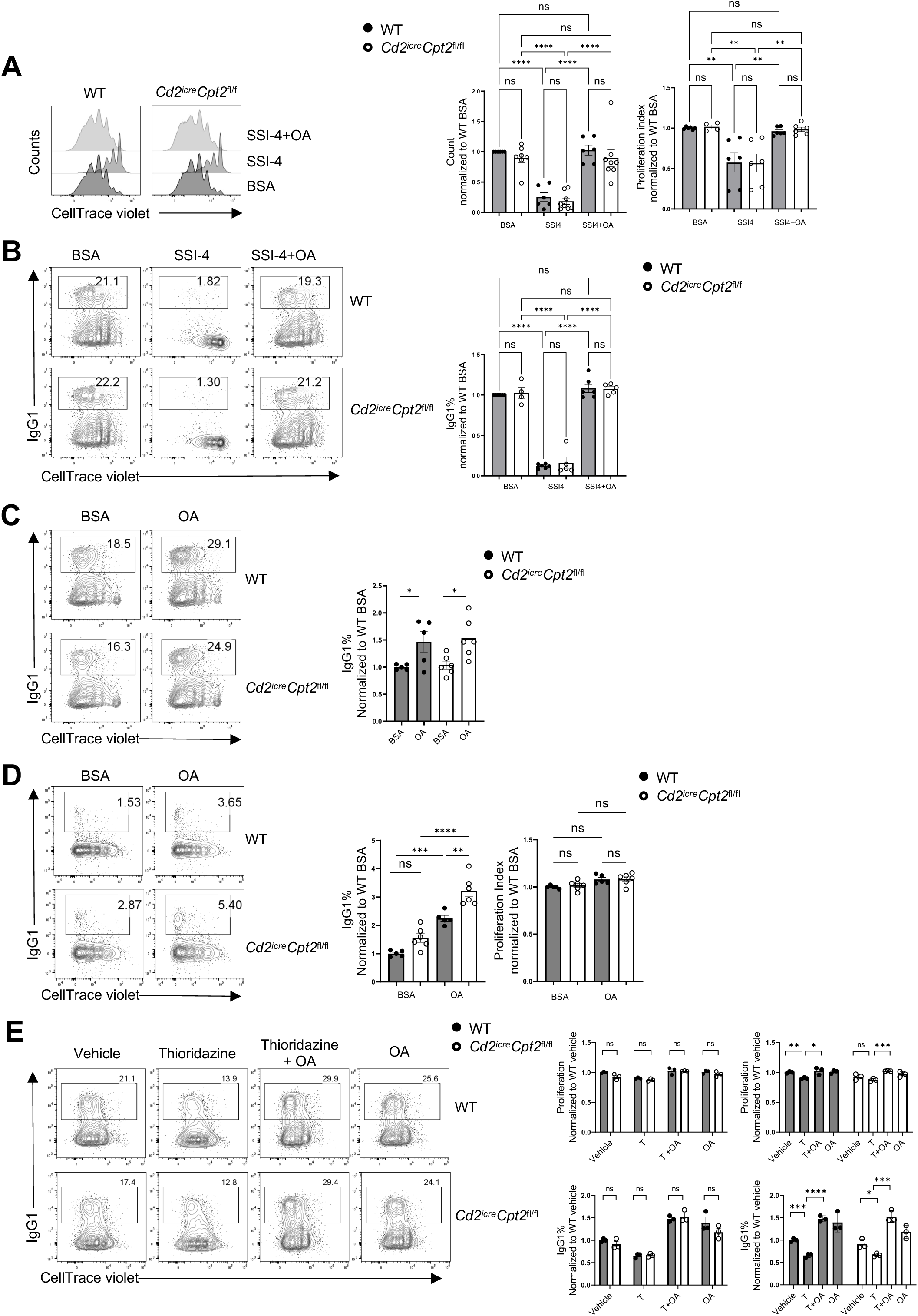
*Cpt2* is dispensable for B cell proliferation, survival, and class switch in vitro. (A) Left, cell proliferation measured by dilution of CellTrace Violet (CTV) dye in B cells from spleens of WT and *Cd2^iCre^Cpt2*^fl/fl^ mice. B cells were stimulated with LPS/IL4/Baff and treated with BSA (50 µM) as control, or SSI-4 (0.1 µM), or SSI-4 with OA (50 µM) for 3 days. Right, summary of cell counts, and proliferation index normalized to WT BSA group. (B) Left, CTV and IgG1 expression on LPS/IL4/Baff activated B cells from WT and *Cd2^iCre^Cpt2*^fl/fl^ mice. Right, summarized IgG1^+^ percentages normalized to WT BSA group. (C) Left, proliferation and IgG1 expression on LPS/IL4/Baff activated B cells from WT and *Cd2^iCre^Cpt2*^fl/fl^ mice treated with BSA (50 µM) as control, or OA (100 µM). Right, IgG1 percentage comparison between two treatments of WT and KO B cells separately. (D) Left, cell proliferation and IgG1 expression on CpG/IL4/IL5 activated B cells from WT and *Cd2^iCre^Cpt2*^fl/fl^ mice treated with BSA (50 µM) as control, or OA (100 µM). Right, summarized IgG1^+^ percentages, and proliferation index normalized to WT BSA group. (E) Left, CTV and IgG1 expression on LPS/IL4/Baff activated B cells from WT and *Cd2^iCre^Cpt2*^fl/fl^ mice treated with vehicle, thioridazine (2.5 µM) with or without OA (100 µM). Right, two formats of summarized proliferation index (above) and IgG1^+^ percentages (below) normalized to WT vehicle group. *p* values were calculated using one-way ANOVA (A, B, and D) or unpaired Student t-test (C and E). ns, not significant, \**p* < 0.05, \*\**p* < 0.01, \*\*\**p* < 0.001, \*\*\*\**p* < 0.0001. Data are representative of at least three (A–E) independent experiments. Error bars represent SEM.

However, no significant differences were noted between the two groups. Additionally, the exogenous OA rectified the reduced proliferation and IgG1 expression induced by thioridazine treatment in both WT and CPT2 deficient B cells (Figure 3E). Taken together, these data indicate that, first, CPT2 deficiency does not affect B cell proliferation and IgG1 expression *in vitro*; second, the effects of exogenous OA on B cells, with or without SCD inhibitor, are independent of CPT2; lastly, peroxisomal FAO may not compensate for CPT2 deficiency in B cells because WT B cells and CPT2 deficient B cells responded to peroxisome inhibitor, with or without exogenous OA, in a similar manner.

### *Cpt2* deletion does not affect NP-OVA immunization induced B cell response *in vivo*

Using *Cpt2* targeting shRNA delivered by retrovirus, a previous study has indicated that CPT2 was critically required for optimal germinal center (GC) formation in response to hapten 4- hydroxy-3-nitrophenylacetyl (NP) immunization (22). Hence, we hypothesized that *Cd2^iCre^Cpt2*^fl/fl^ mice would have a reduced GC formation and overall humoral immune responses, which might be reflected by reduced immunoglobulin level at baseline and/or after immune challenges. We first measured the serum immunoglobulin levels at baseline. WT and *Cd2^iCre^Cpt2*^fl/fl^ mice had equivalent serum concentrations of all immunoglobulin isotypes measured, including IgM, IgG1, IgG2a, IgG2b, IgG3, and IgA (Figure 4A). Then we immunized mice with NP-OVA precipitated in alum. Generation of GC B cells and NP-specific GC B cells were not affected by CPT2 deficiency (Figure 4B, 4C). *Cd2^iCre^Cpt2*^fl/fl^ mice also exhibited normal plasmablast formation after NP-OVA immunization (Figure 4D). Moreover, *Cpt2* deletion did not affect differentiation of follicular helper T (Tfh) cells (Figure 4E). Consistent with the normal GC and plasmablast formation, the production of all affinity (anti-NP23) and low affinity (anti-NP2) immunoglobulins was not significantly altered in *Cd2^iCre^Cpt2*^fl/fl^ mice (Figure 4F). Thus, our data indicates that CPT2 expression in lymphocytes is dispensable for NP- OVA immunization elicited humoral immune responses.

**FIGURE 4.**
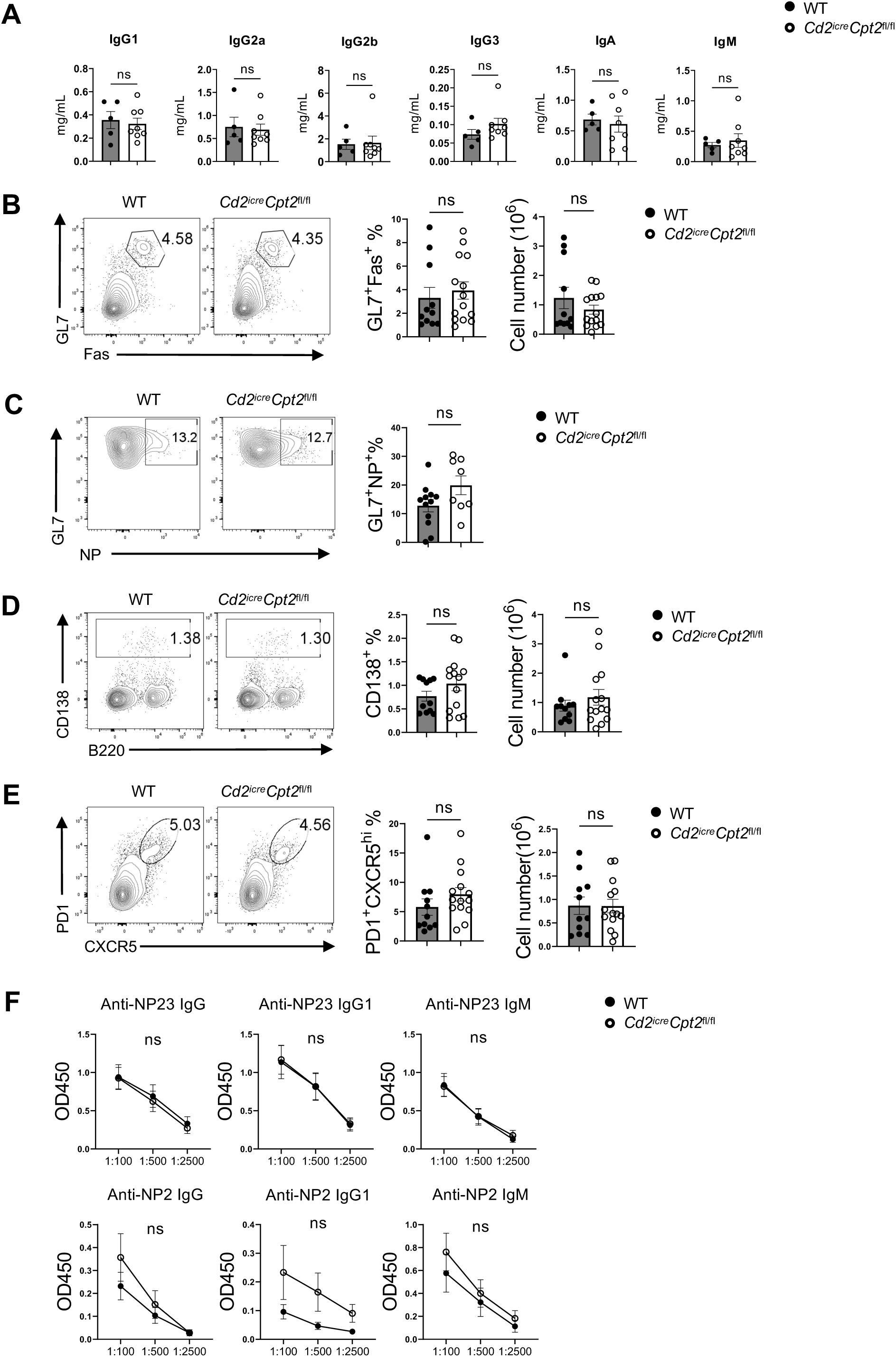
*Cpt2* deletion may not affect NP-OVA immunization induced B cell response *in vivo*. (A) The baseline levels of immunoglobulin (Ig) G1, IgG2a, IgG2b, IgG3, IgA and IgM in mouse sera collected from WT and *Cd2^iCre^Cpt2*^fl/fl^ mice. (B-E) WT and *Cd2^iCre^Cpt2*^fl/fl^ mice were immunized with NP-OVA and analyzed after 14 days. Left, flow cytometry of GC expression in B220^+^ cells (B), GL7 and NP expression in GC (C), B220^int^CD138^+^ expression (D), and expression of PD-1 and CXCR5 among CD4^+^ T cells (E) from spleen lymphocytes. Right, the summaries of GC B cells (B), GL7^+^NP^+^ populations (C), B220^int^CD138^+^ plasmablasts (D), and PD-1^+^CXCR5^hi^ Tfh cells (E). (F) ELISA test of serum anti-NP23 and anti-NP2 immunoglobulins from NP-OVA immunized mice presented as absorbance at 450 nM (A_450_). *p* values were calculated using Student’s t test (A–E). ns, not significant. Data are representative of three (B-F) independent experiments. Error bars represent SEM.

### B cell intrinsic CPT2 is dispensable for NP-OVA induced humoral immunity

To address whether B cell intrinsic CPT2 expression is necessary for humoral immunity *in vivo*, we generated mixed chimera mice. We mixed BM cells from WT or *Cd2^iCre^Cpt2*^fl/fl^ mice with CD45.1 congenic mouse BM cells at a ratio of 1:1. The mixed BM cells were transferred into *Rag1^−/−^* mice that lack mature T and B cells. In such mixed chimera mice, WT and CPT2 deficient B cells would receive help from WT T cells. Ten weeks after BM reconstitution, the chimera mice were immunized with NP-OVA. We observed equivalent GC, NP-specific GC, plasmablast and Tfh formation in the *Cd2^icre^Cpt2*^fl/fl^ chimera mice relative to WT chimera mice (Figure 5A-5D). The production of NP specific antibodies was also largely normal (Figure 5E). Thus, B cell intrinsic CPT2 expression is not necessary for normal GC and plasmablast formation in response to NP-OVA immunization.

**FIGURE 5.**
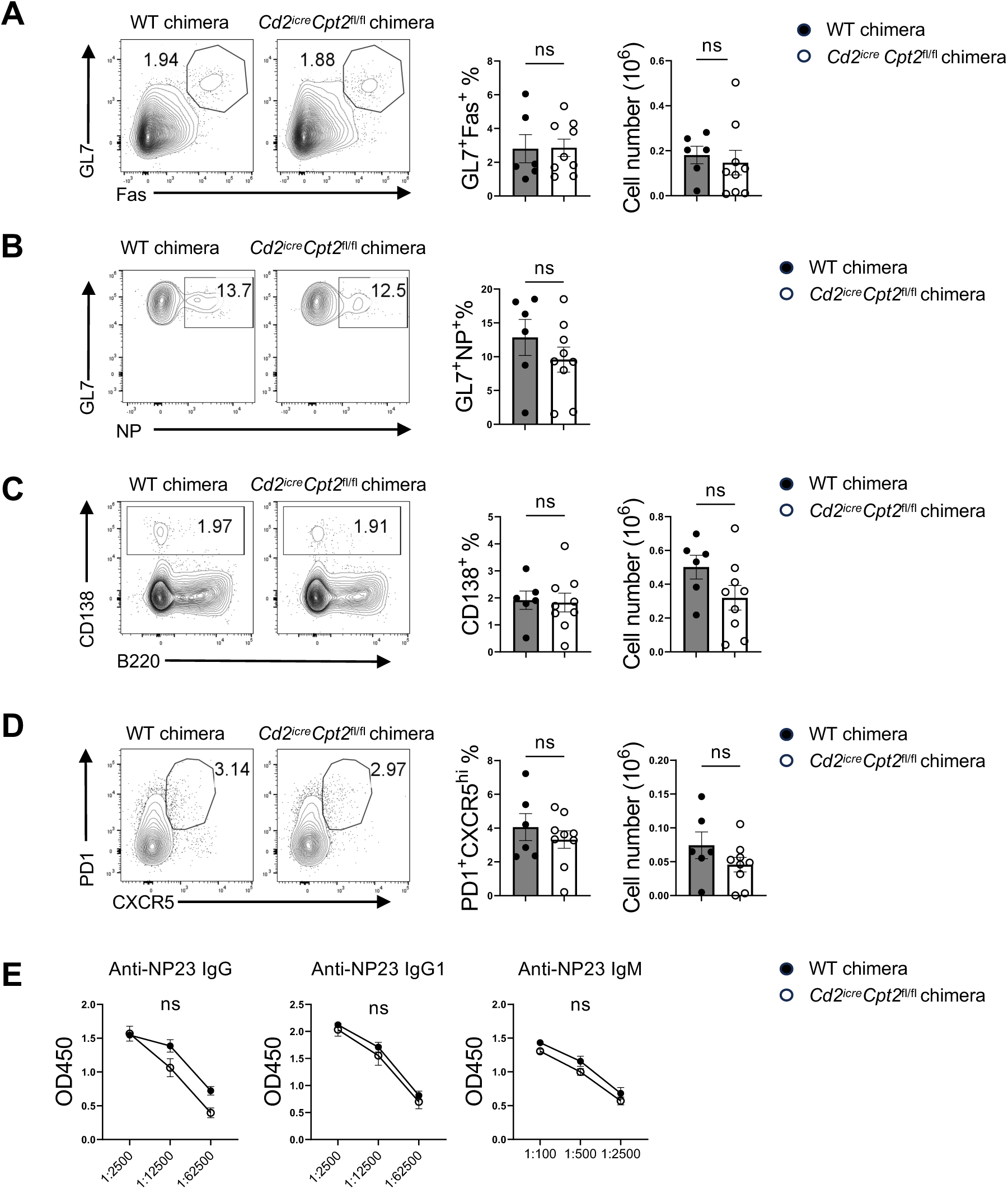
B cell intrinsic CPT2 expression is not required for humoral immunity *in vivo*. (A-D) WT and *Cd2^iCre^Cpt2*^fl/fl^ chimera mice were immunized with NP-OVA and analyzed after 14 days. Left, flow cytometry of GC expression in B220^+^ cells (A), GL7 and NP expression in GC (B), B220^int^CD138^+^ expression (C), and expression of PD-1 and CXCR5 among CD4^+^ T cells (D) from spleen lymphocytes. Right, the frequencies of GC B cells (A), GL7^+^NP^+^ populations (B), B220^int^CD138^+^ plasmablasts (C), and PD-1^+^CXCR5^hi^ Tfh cells (D). (E) Serum anti-NP23 and anti-NP2 immunoglobulins from NP-OVA immunized mice were measured by ELISA. *p* values were calculated using Student’s t test (A–D). ns, not significant. Data are representative of two (A-E) independent experiments. Error bars represent SEM.

### CPT2 is not required for humoral immunity against influenza infection

B cells play a critical role in host defense against influenza infection (23). To evaluate if CPT2 expression in lymphocytes contributes to anti-viral humoral immunity, we infected WT and *Cd2^icre^Cpt2*^fl/fl^ mice with H1N1 influenza A virus intranasally. *Cd2^icre^Cpt2*^fl/fl^ mice showed a similar weight decrease as WT after influenza infection (Figure 6A). We examined GC and plasmablast formation in spleens and mediastinal lymph nodes (Figure 6B, 6C) and found equivalent responses between WT and *Cd2^icre^Cpt2*^fl/fl^ mice. Tfh cells also differentiated normally in the absence of Cpt2 (Figure 6D). Finally, deletion of *Cpt2* did not affect the production of antibodies against influenza (Figure 6E). Therefore, CPT2 is dispensable for humoral immunity upon respiratory viral infection.

**FIGURE 6.**
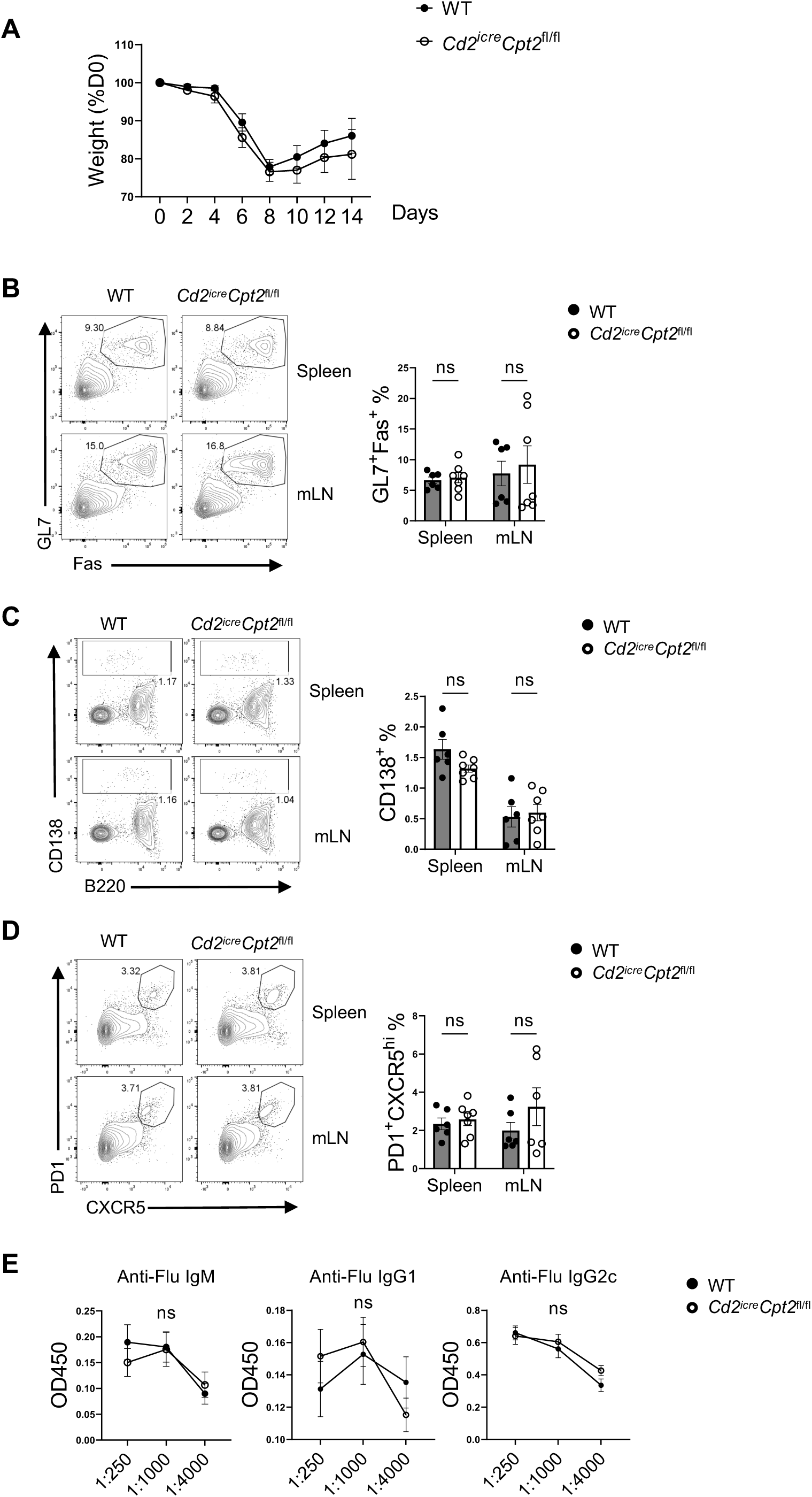
*Cpt2* is not required for humoral immunity against influenza infection. (A) Weight change of WT and *Cd2^iCre^ Cpt2*^fl/fl^ mice following influenza infection. (B-D) Left, flow cytometry analysis of GC B cells (B), B220^int^CD138^+^ expression (C), and expression of PD-1 and CXCR5 among CD4^+^ T cells (D) from spleens and mediastinal lymph nodes at day 14 following infection. Right, the frequencies of GC B cells (B), B220^int^CD138^+^ plasmablasts (C), and PD-1^+^CXCR5^hi^ Tfh cells (D). (E) Influenza virus-specific antibodies IgM, IgG1, IgG2c in sera were measured using ELISA. *p* values were calculated using Student’s t test (A–E). ns, not significant. Data are representative of two (B-E) independent experiments. Error bars represent SEM.

### CPT2 deficiency does not impair the immune response to TNP-LPS immunization

B cells can respond to thymus-dependent (TD) antigens, including NP-OVA and influenza virus, and thymus-independent (TI) antigens, such as TNP-LPS. Our data so far indicate that CPT2 is dispensable for humoral immune response to TD antigens. We next tested whether CPT2 deficiency affected TI response by immunizing WT and *Cd2^icre^Cpt2*^fl/fl^ mice with TNP-LPS. We found that CPT2 deficiency did not affect plasmablast formation (Figure 7A) or B cell activation measured by GL7 and IgD expression (Figure 7B). Importantly, *Cd2^icre^Cpt2*^fl/fl^ mice produced the same level of anti-TNP IgM and IgG antibodies as WT mice did (Figure 7C). Therefore, our data indicates that CPT2 is dispensable for humoral responses to TI antigen TNP-LPS.

**FIGURE 7.**
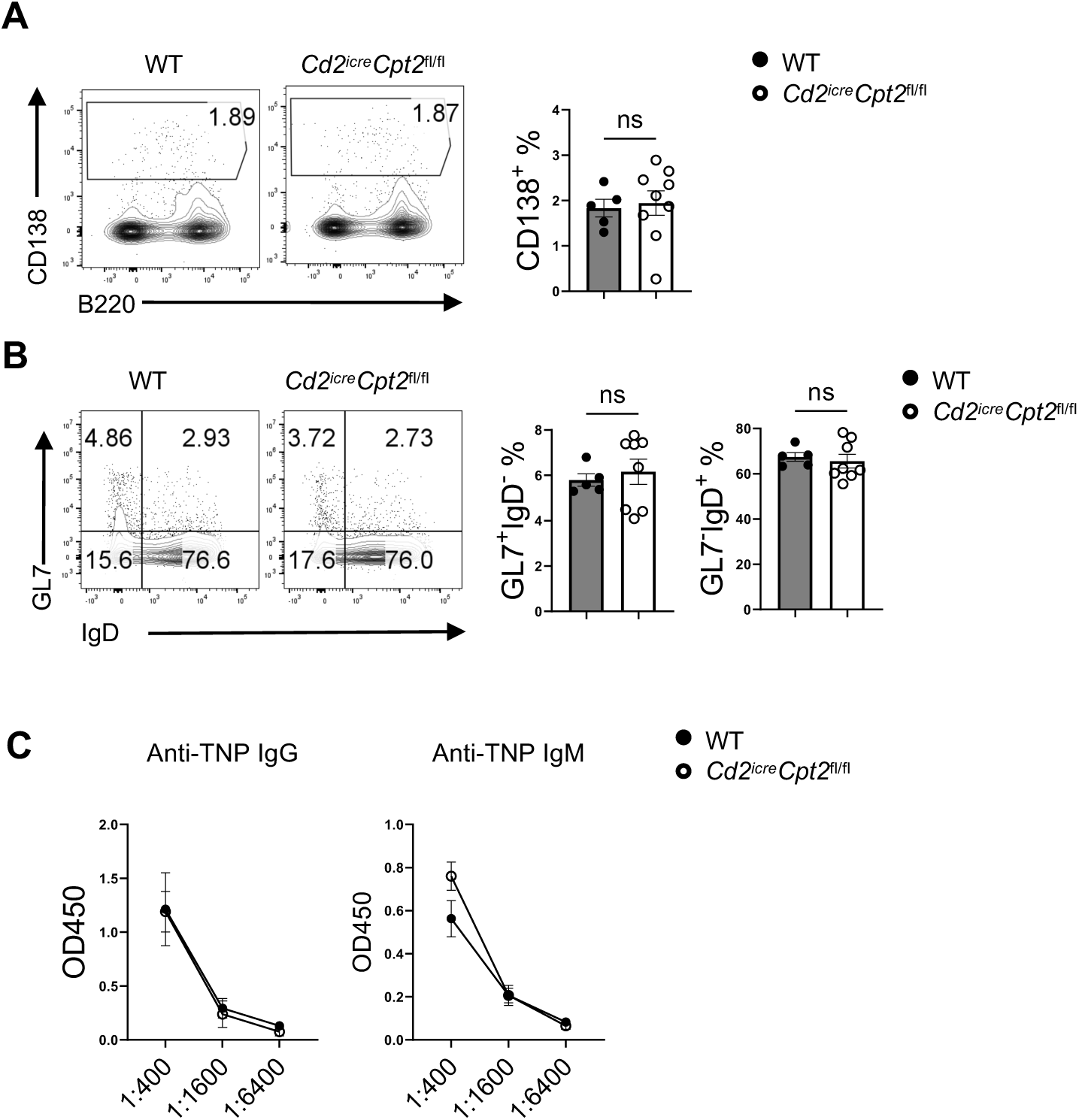
*Cpt2* deletion does not impact the immune response to TNP-LPS immunization. (A-B) WT and *Cd2^iCre^ Cpt2*^fl/fl^ mice were immunized with TNP-LPS and analyzed after 14 days. Left, flow cytometry of B220^int^CD138^+^ expression (A), and expression of GL7 and IgD among B220^+^ cells (B) from spleen lymphocytes. Right, the frequencies of B220^int^CD138^+^ plasmablasts (A), and GL7^+^IgD^-^ and GL7^+^IgD^-^ cell populations (B). (C) Anti-TNP IgG and IgM in sera measured by ELISA. *p* values were calculated using Student’s t test (A–B). ns, not significant. Data are from one (A-C) experiment. Error bars represent SEM.

## DISCUSSION

Metabolic plasticity is characteristic of immune cells. Recent stable isotope tracing experiments have demonstrated that T cells can utilize glucose, lactose, glutamine and acetate to fuel their biosynthetic and bioenergetic needs (24–27). Fatty acid is a major energy source. LCFA beta- oxidation in mitochondrial has been shown to be critical for the generation and maintenance of memory T cells, Tregs and GC B cells, as well as NK cell effector function (17, 22, 28, 29). Yet, contradictory findings have been presented because genetic ablation of CPT1A does not affect the differentiation of memory T cells and Tregs (16). Thus, it remains contentious whether LCFA beta-oxidation is critically required for lymphocyte differentiation and function. Our study was undertaken to provide a definitive answer to the dependency of CPT mediated LCFA beta- oxidation for humoral immunity.

Using a mouse line with lymphocyte specific deletion of CPT2, the obligatory enzyme that mediates LCFA transfer across the mitochondrial inner membrane for beta-oxidation, we demonstrated that CPT2 is necessary for optimal LCFA oxidation and mitochondrial metabolism in B cells. Yet, despite highly reduced LCFA beta-oxidation, B cell activation and proliferation *in vitro* and humoral immune responses towards both TD and TI antigens, including GC and plasmablast formation, and antibody production, were not significantly affected by CPT2 deficiency *in vivo.* Thus, our data indicate that CPT2 mediated LCFA beta-oxidation is largely dispensable for humoral immunity. It is currently unclear why our data contradict the earlier study, which showed a reduced GC formation when *Cpt2* was knockdown in B cells (22). There are several potential explanations for such divergent observations. First, different gene targeting approaches were utilized, Cre-Loxp mediated genetic deletion in our study versus retroviral introduction of *Cpt2* targeting ShRNA in the prior work. Second, different B cell responses were evaluated, polyclonal reaction in our study versus monoclonal reaction with B1-8 transgenic B cells in the prior work. Nevertheless, because we did not detect any significant changes in terms of antibody production, it seems unlikely that CPT2 ablation has a substantial impact on humoral immunity. Therefore, our data suggest that other energy sources could fully compensate for the loss of LCFA beta-oxidation for B cell response *in vivo,* again highlighting the metabolic flexibility of B cells. It remains to be seen whether CPT2 is required for the differentiation of memory B cells in future studies.

Fatty acids and their derivatives can in theory affect immune cells through mechanisms other than mitochondrial beta-oxidation. For example, they are an integral part of cellular and organelle membranes (30). Monounsaturated fatty acids can help control endoplasmic reticulum stress (21, 31). Finally, LCFA can be oxidized in peroxisomes independent of CPT2 (22, 32). Our data indicated that peroxisomal oxidation likely does not compensate for the loss of CPT2.

Therefore, our results suggest that loss of LCFA beta-oxidation can be fully compensated by hitherto undefined mechanisms in B cells, which warrant further investigation.

## ACKNOWLEDGEMENTS

We thank Dr. Jie Sun for sharing research reagents. We acknowledge the Microscopy and Cell Analysis Core facility at Mayo Clinic Rochester. We acknowledge the NIH (grants R01 AI 162678 and R01 AR077518) for supporting this work in H.Z.’s laboratory, and R01 CA225680 to T.H.’s laboratory.

## AUTHOR CONTRIBUTIONS

M.L. and H.Z. conceived the project, designed the research, interpreted the data and wrote the manuscript. M.L. and X.Z. prepared the materials and carried out the experiments. X.X.Z. performed the influenza infection. Yanfeng Li managed the mouse colony and performed the molecular biology experiments. Yuzhen Li provided research material. T.H. performed the stable isotope labeled FAO assay.

